# Machine learning based detection of genetic and drug class variant impact on functionally conserved protein binding dynamics

**DOI:** 10.1101/724211

**Authors:** Gregory A. Babbitt, Ernest P. Fokoue, Joshua R. Evans, Kyle I. Diller, Lily E. Adams

## Abstract

The application of statistical methods to comparatively framed questions about protein dynamics can potentially enable investigations of biomolecular function beyond the current sequence and structural methods in bioinformatics. However, chaotic behavior in single protein trajectories requires statistical inference be obtained from large ensembles of molecular dynamic (MD) simulations representing the comparative functional states of a given protein. Meaningful interpretation of such a complex form of big data poses serious challenges to users of MD. Here, we announce DROIDS v3.0, a molecular dynamic (MD) method + software package for comparative protein dynamics, incorporating many new features including maxDemon v1.0, a multi-method machine learning application that trains on large ensemble comparisons of concerted protein motions in opposing functional states and deploys learned classifications of these states onto newly generated protein dynamic simulations. Local canonical correlations in learning patterns generated from self-similar MD runs are used to identify regions of functionally conserved protein dynamics. Subsequent impacts of genetic and drug class variants on conserved dynamics can also be analyzed by deploying the classifiers on variant MD runs and quantifying how often these altered protein systems display the opposing functional states. Here, we present several case studies of complex changes in functional protein dynamics caused by temperature, genetic mutation, and binding interaction with nucleic acids and small molecules. We studied the impact of genetic variation on functionally conserved protein dynamics in ubiquitin and TATA binding protein and demonstrate that our learning algorithm can properly identify regions of conserved dynamics. We also report impacts to dynamics that correspond well with predicted disruptive effects of a variety of genetic mutations. In addition, we studied the impact of drug class variation on the ATP binding region of Hsp90, similarly identifying conserved dynamics and impacts that rank accordingly with how closely various Hsp90 inhibitors mimic natural ATP binding.

**Statement of significance:** We propose a statistical method as well as offer a user-friendly graphical interfaced software pipeline for comparing simulations of the complex motions (i.e. dynamics) of proteins in different functional states. We also provide both method and software to apply artificial intelligence (i.e. machine learning methods) that enable the computer to recognize complex functional differences in protein dynamics on new simulations and report them to the user. This method can identify dynamics important for protein function, as well as to quantify how the motions of molecular variants differ from these important functional dynamic states. For the first time, this method of analysis allows the impacts of different genetic backgrounds or drug classes to be examined within the context of functional motions of the specific protein system under investigation.

## Introduction

The physicist Richard Feynman is said to have once famously quipped, ‘all biology is ultimately due to the wiggling and jiggling of atoms’. Stated with more precision, Feynman’s conjecture would imply that all biological function can ultimately be understood by analyzing rapid molecular motions in biomolecular structures as they alter or shift their functional state(s). Many decades later, these functional shifts in molecular dynamics are being illuminated by structural and computational biology. Examples of functionally altered dynamics include the destabilization of inter-residue contacts, in both disease malfunction and normal signal activation, as well as the stabilization of inter-residue contacts during protein folding, the formation of larger complexes, and various other binding interactions to small molecules. And while the functional role of rapid vibrations revealed by short term molecular dynamic (MD) simulations has been debated in the past, more recent empirical and computational studies have clearly demonstrated that differences in both rapid and directed vibrations can drive longer term functional conformational change (1, 2). From a broader perspective, if Feynman’s conjecture is true, then the specific details of a given protein system’s biomolecular dynamics will represent a potentially large source of latent variability in our functional understanding of the genome; a problem largely ignored by those disciplines currently generating the vast amounts of static forms of ‘omic’ type data (i.e. DNA sequence, transcript level, and protein structure)(3). However, in the last decade, simultaneous advances in the development of graphics hardware and biomolecular force fields has elevated our ability to computationally simulate MD long enough to capture ns to μs timescales for moderately-sized proteins (4, 5), and finally ‘see’ some of their functionally relevant motions. And now, the application of proper statistical comparisons of ensembles of short-timed framed MD simulation can potentially enable meaningful interpretations of comparative questions about protein dynamics (6). But due to the richly complex structure of data underlying the moving images generated by MD software, functional interpretation of modern MD simulations poses a serious challenge to current users, especially with comparatively-framed questions, where large ensembles of many production runs need to be generated and subsequently analyzed. A potential solution to this problem exists with the application of machine learning to the feature extraction and classification of the dynamic differences between ensembles of MD runs. These ensembles can be designed to represent pair-wise functional states of biomolecular systems (e.g. before/after chemical mutation or binding). Therefore, the high performance accelerated computation used to generate simulated protein motions for comparison can be effectively partnered with high performance methods for optimally extracting and learning the underlying dynamic feature differences defining the different functional states of proteins. Although machine learning has recently been applied to individual MD studies for a variety of specific tasks (7–9), there is no current software platform for the general application of machine learning to comparative protein dynamics.

In 2018, we released DROIDS v1.2 and v2.0 (<u>D</u>etecting <u>R</u>elative <u>O</u>utlier <u>I</u>mpacts from molecular <u>D</u>ynamic <u>S</u>imulation), a GPU accelerated software pipeline designed for calculating and visualizing statistical comparisons of protein dynamics drawn from large repeated ensembles of short dynamic simulations representing two protein states (6). This application allowed simple visual and statistical comparison of protein MD ensembles set up in any way the user wanted to define them. Here, we announce the release of DROIDS v3.0, which now offers multiple pipelines tailored for specific functional comparisons of systems comprised of combinations of proteins, nucleic acids, and small ligand molecules. Comparisons can include different temperatures, different protein binding states (i.e. to DNA, drugs, toxins or natural ligands), or divergent genetic/epigenetic mutant states. We also include a major new machine learning tool, maxDemon v1.0, a multi-machine learning post-processing application for DROIDS that trains on the data representing the comparatively divergent functional dynamic states, and subsequently identifies functionally conserved dynamics and genetic and/or drug class binding variant effects when deployed on new MD simulations representing these variants of interest. Thus, much like James Clerk Maxwell’s mythical creature (10), maxDemon derives important information from all atom resolution observation of dynamic motion. The three primary features/aims of our newly expanded software is to (A) improve user experience in comparative protein dynamics, (B) enable the local detection of functionally conserved protein dynamics, and to (C) enable the assessment of the local dynamic impacts of both genetic and drug class variants within the functional context of protein system of interest. Because the machine learning model we employ is trained on MD data representing normal functioning dynamic states of a protein, this metric of impact is highly context dependent to how a given mutation or drug impacts a specific protein. Thus, it potentially gives considerably more functional relevance to the analysis of variants when compared to more general database-derived metrics of mutational tolerance (e.g. SIFT, PolyPhen2 etc.). In Table 1, we list five primary methodological pipelines in available in DROIDS 3.0+maxDemon to address functional questions in comparative protein dynamics. In our results and discussion here, we present data on four case studies of functional protein dynamics that include feature extraction and classification of (A) a simple temperature shift in ubiquitin dynamics, (B) mutational impacts on ubiquitin binding dynamics, (C) mutation specific impacts on DNA binding of TATA binding protein, and (D) comparison of binding dynamics of drug class variants that mimic ATP binding in Hsp90.

**Table 1.** Common learner assisted comparative protein dynamic investigations enabled by DROIDS 3.0 + maxDemon 1.0.

| QUESTION | DROIDS 3.0 training comparison | Deployment of learners in maxDemon | Important notes |
| --- | --- | --- | --- |
| Measure dynamic tolerances of single protein to various genetic mutations | Two sets (ensembles) of MD on the same protein at the same temperature | MD run on one or more genetic mutant structures | Isolates MD impacts of mutation(s) from natural variability in self-similar dynamics |
| Measure dynamic tolerances of DNA binding interaction to genetic mutation(s) | MD ensembles comparing both the unbound and DNA bound protein | MD run on one or more unbound genetic mutant structures | Isolates MD impacts of mutation from natural binding function of the system |
| Measure dynamic tolerances of individual genetic differences to a given drug | MD ensembles comparing both the unbound and drug bound protein | MD run on one or more drug-bound genetic mutant structures | Isolates MD impacts of mutation from novel drug binding function of the system |
| Measure dynamic similarities of different drug candidates to natural ligand binding interaction | MD ensembles comparing both the unbound and ligand bound protein | MD run on one or more drug variant bound structures | Isolates MD impacts of drug candidates from the natural binding function of the ligand |
| Measure evolution of novel dynamics in paralog genes | MD ensembles comparing two ortholog proteins (i.e. same gene different species) | MD runs on one or more paralogs (i.e. duplicated genes in same species) | Isolates potential MD novelty in duplicated gene product from nonfunctional or neutral changes in different species |

## Materials and Methods

### Overview of comparative dynamics and visualization with DROIDS v3.0

Our DROIDS method/software leverages several important key concepts when making comparisons between MD runs. The method utilizes structural alignment to restrict comparison of dynamics between individual homologous amino acids. The method also restricts comparison averaged over atoms common to all amino acids (i.e. backbone C, N, O and C_α_). The method also employs statistical ensembling to make a robust comparison between protein dynamics in different functional states (6). While this is computationally intensive, it is necessary because of the inherent chaotic nature and unpredictability of single protein trajectory projections. This logic is analogous to the many storm tracks repeatedly modeled by meteorologists to gain statistical confidence in a hurricane weather forecast, where an ensemble of model runs all with slightly different initial conditions has far more predictive power than any single simulation. In DROIDS, the user can decide how large the MD ensembles need to be based upon the inherent stability of the protein under investigation. Generally, an ensemble size of 200 to 300 MD runs at 0.5-1 ns will suffice for most proteins. The dynamics is summarized by calculation of root mean square fluctuations (*rmsf*) over constant time intervals represented by a constant number of image frames defined by the user (thus allowing *rmsf* values to be sampled on repeatedly on an identical and thus comparable scale). The default number of frames (*i*) in the software for a given time slice is n=50 representing 0.01 ns of simulation time. The rmsf value is thus

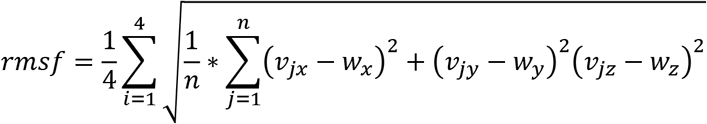

where *v* represents the set of XYZ atom coordinates for *i* backbone atoms (C, N, O, and C_α_) for a given amino acid residue over *j* time points and *w* represents the reference coordinate structure at the beginning of each MD production run for a given ensemble. Therefore, *rmsf* values as defined here represent molecular dynamics at the resolution of a single amino acid backbone segment, and the same resolution at which fine scale protein-level molecular evolution operates via amino acid replacement, insertion and deletion. The *rmsf* is also the most basic underlying functional quantity to extract from MD simulation as its underpins all hierarchical levels of motion (1). Two ensembles of *rmsf* values (a query set and a reference set) are compared to calculate average delta *rmsf* or *dRMSF*. The user can choose to see the average angstrom difference between sets of values, or more preferably the user can calculate the symmetric Kullback-Leibler divergence (i.e. relative entropy) between the two empirical statistical distributions of *rmsf*. The KL divergence generally provides a richer more informative view of dynamic differences with less loss of information than simple averaging. Thus *dRMSF* comparing *rmsf* values for two ensembles of size *m* for a given amino acid is

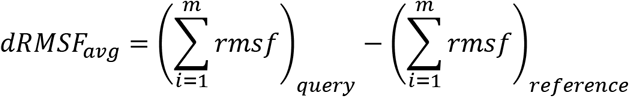

or

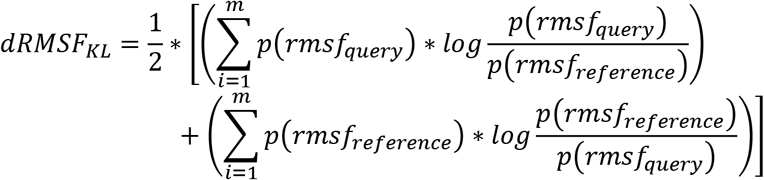

The resulting dRMSF values are color mapped to either still structures or movie images of the dynamics according to either a ‘temperature’ scale where (+) dRMSF = amplified vibration is red and (−) dRMSF = dampened vibration is blue. A ‘stoplight’ scale where (+) dRMSF is green and (−) dRMSF is red is also available. On both scales, neutral values are shaded towards white.

### Functional classification of new MD simulation with maxDemon v1.0

While users can easily employ DROIDS 3.0 to examine ensemble differences between functional genetic or binding states, the application of this knowledge to new MD simulation is nearly impossible due to the inherent complexity of the moving protein behavior. Our new post-processing software, maxDemon 1. 0, uses machine learning to label or classify the differences learned by a previous DROIDS query/reference state comparison when subsequently applied to one or more new MD runs. The machine learning-based detection of variant impacts on functional protein dynamics presented here is outlined schematically in Figure 1. Similar to the statistics for comparative dynamics, the learning algorithms are also applied individually to each amino acid backbone’s ensemble of *rmsf* values. This allows for similar single residue resolution in the results. Learners are also applied within the same user defined time slices of *rmsf* allowing for visualization of time resolution of classification of functional dynamic behaviors as well. The learning performance is summarized by tallying the average classification over all time slices for each amino acid. Thus, an average performance of 0.5 would indicate that the learners are not finding the functional states defined by and trained by the initial DROIDS comparative analysis. Canonical correlations in the positional performance plots are key in detecting sequence encoded functionally conserved dynamics regions, as well as genetic and drug class variant impacts to these functional regions as well. This is described with more formality below.

**Figure 1.**
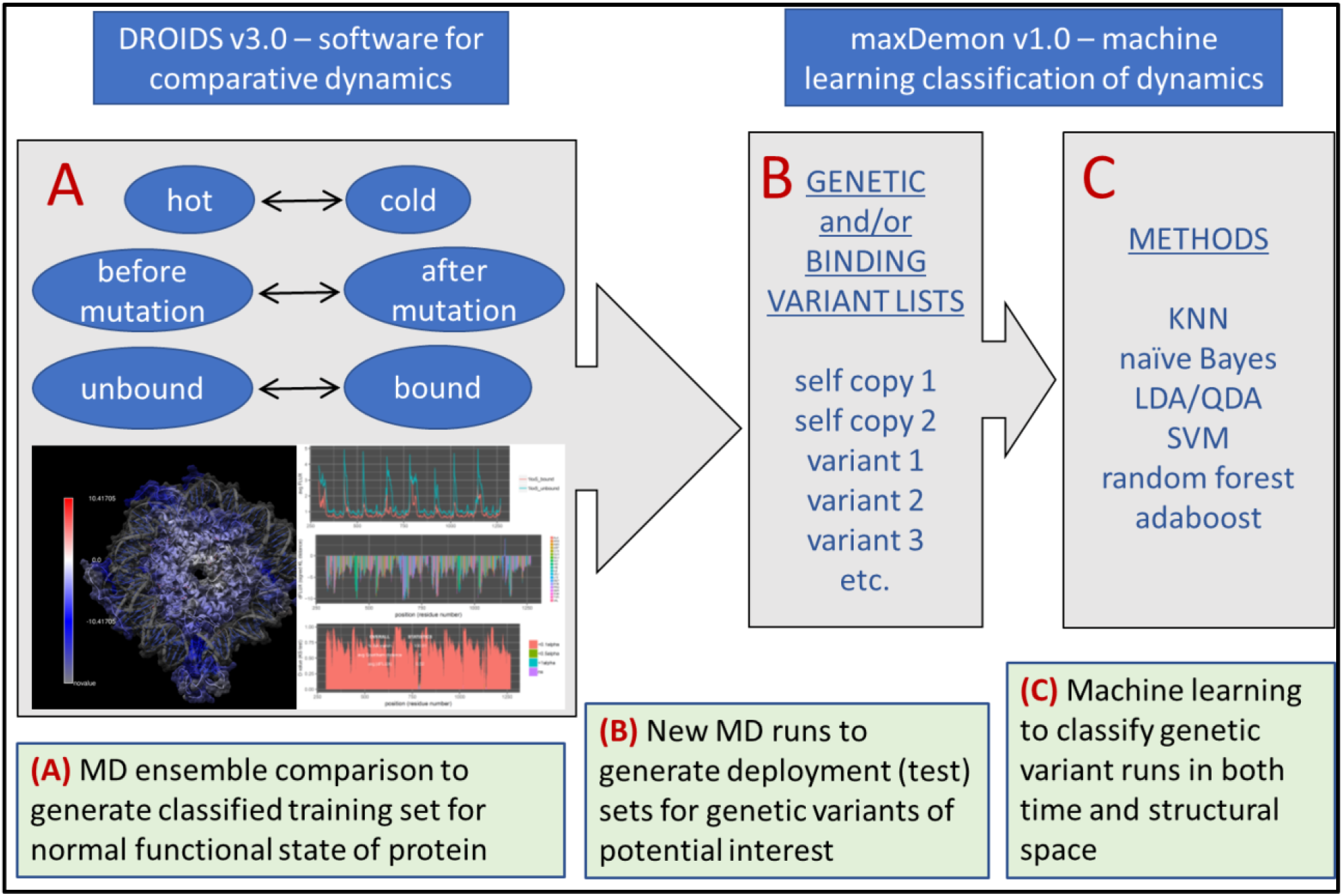
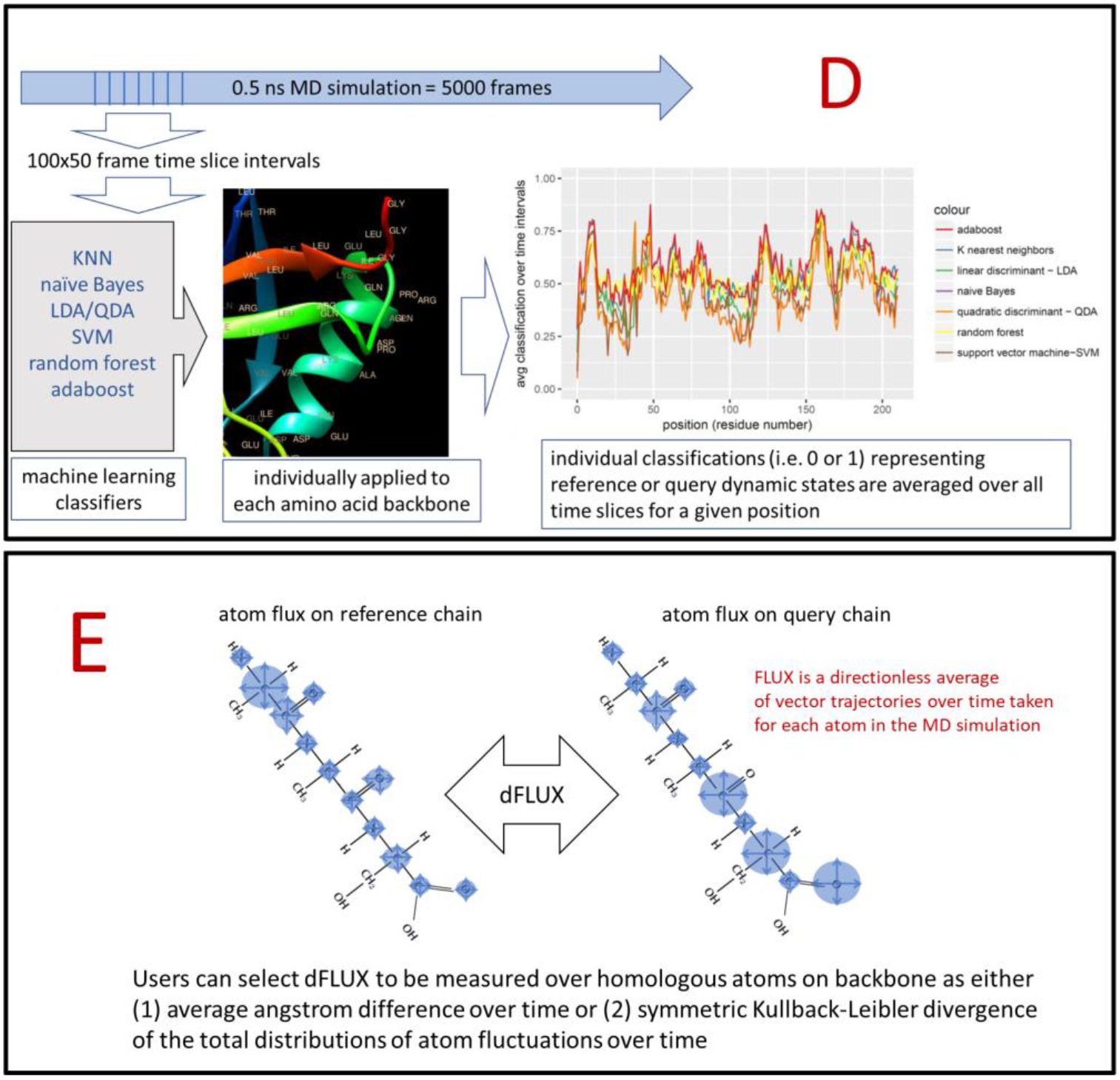
Schematic overview of DROIDS 3.0 + maxDemon 1.0 software for machine learning-based detection of variant impacts on functionally conserved protein dynamics. The pipeline starts with (A) generation of two large ensembles of molecular dynamic (MD) simulations that represent a functional comparison of protein states (e.g. mutation, binding or environmental change). The root mean square fluctuations (rmsf) of protein backbone atoms in these ensembles are comparatively analyzed/visualized (i.e. using DROIDS) and are also later used as pre-classified training data sets for machine learning (i.e. using maxDemon). Note: in the pictured DROIDS analysis of nucleosome shows overall dampening of rmsf in the histone core with maximal dampening where the histone tails cross the DNA helix (B) New MD simulations are generated on two structures self-similar to the query state of training as well as a list of functional variants, and (C) up to seven machine learning methods are employed to classify the MD in the self-similar and variant runs according to the functional comparison defined by the initial training step. (D) The performance of learning is defined by average value of classification (i.e. 0 or 1) over 50 frame time slices for each amino acid position and regions of functionally conserved dynamics are later identified by significant canonical correlations in this learning efficiency (i.e. Wilk’s lamda) in self-similar MD runs. The impacts of variants are defined by relative entropy of variant MD compared to the MD in the self-similar runs and plotted when this entropy is significantly different from the variation in self-similarity (i.e. bootstrapped z-test). (E) A visual representation of the difference in local rmsf (dFLUX) is typically calculated using symmetric Kullback-Leibler (KL) divergence between the two distributions of rmsf in the training MD ensembles.

### Machine learning training and validation

The feature vectors (*X*) for machine learning are collections of *rmsf* values (x_i_) labeled according to a query (q) and reference state (r) defined by the DROIDS MD comparison (i.e. where labels y_i_ are *q = 1* and *r* = 0).

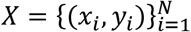

The length of the vector (N) is defined by the length of the MD production run chosen by the user and the size of the ensemble of MD production runs taken. Thus if the user chooses an ensemble of 200 MD production runs each at a time length of 0.5 ns (= 2500 frames) and uses the default time interval of 50 frames to calculate any given interval of *rmsf* then the resulting feature vector will contain 20,000 data values for training (i.e. 10,000 values each for *q* and *r*).

Users create a ‘stacked model’ or meta-model containing up to seven different machine learning classification algorithms including K-nearest neighbors, naïve Bayes, linear discriminant analysis, quadratic discriminant analysis, random forest, adaptive boosting and support vector machine (with kernel options including parameter tuned linear, polynomial, laplace and radial basis functions). R packages employed here are KNN, MASS, kernlab, randomForest and ada. We restricted machine learning to ‘shallow’ learning methods due to the relatively small datasets created when resolving dynamics of protein systems to short slices of time over single amino acids and also because of the robustness of the R packages when applied sequentially over time and structural space. Therefore, we do not yet support implementation of deep learning neural networks. For methodologically robust results on small proteins, we generally recommend users select all seven available methods. As real features of dynamics should be detectable by any method of learning, the agreement of classification obtained by the creation of a stacked model utilizing different learning methods makes the learning less sensitive to methodological artifacts. Depending upon system resources, users can choose to include or omit methods from four categories of learning (i.e. instance-based = KNN, probabilistic = NB / LDA / QDA, black box = SVM, and ensemble learning = randomForest / adaboost. Users will want to use as many as their system resources can handle, however for faster processing, a minimum of three of the seven learning methods can be chosen. Currently, most methods run on single CPU cores, however, the more CPU intensive methods of random forest and adaboost algorithms are programmed to use all available CPU cores found on the system. SVM is often the slowest method for larger protein systems and can be omitted when more than 300 residues are present in the protein simulation.

After learners are trained on the query and reference ensembles, they are validated on a new MD run that matches the state of the reference MD runs during training. For example, when analyzing a binding interaction where the reference ensemble of training runs are conducted in the unbound protein state, a new run will be conducted in the unbound state and a line plot of the machine learning performance (i.e. precision, recall and accuracy) will be generated for all positions on the protein. It would be expected that if comparative differences in dynamics observed in the training set have a genuine relation to function(s) defined during training, they will display repeated behavior in the new reference run and be identified by the stacked learning model generating local peaks in learning performance (i.e. accuracy) at functional regions (Figure 1D). Learner performance for a given machine learning method is defined as

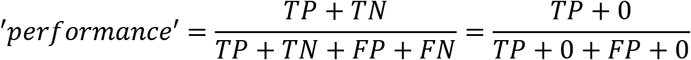

Where TP, TN, FP, and FN are true positive, true negative, false positive, and false negative classifications resp. The zero value terms arise because the validation is conducted on simulation representing just the reference state of the DROIDS comparison (where y_i_ = 0). Therefore, accuracy, and precision are algebraically collapsed to a single equivalent performance metric while recall is always equal to 1.

### Identifying regions of conserved dynamics

Functionally conserved dynamics are defined as ‘repeated or self-similar and sequence-dependent dynamics’ discovered after training machine learners on the functional state ensembles derived with DROIDS. Conserved dynamics are detected via significant canonical correlations in position specific learning performance patterns after the deployment of learners on new MD simulation runs that were setup identically to the reference dynamic state defined by the MD ensemble training set. We expect that functionally conserved dynamics will be sequence encoded and therefore should display a repeated position dependent signature in our learned pattern profiles whenever MD runs are set up identically to MD upon which learners were trained. Therefore, a significant local canonical correlation (i.e. Wilk’s lambda) between learning performance profiles of self-similar MD runs can be used to detect local regions of conserved protein dynamics.

To detect functionally conserved dynamics after training and validation, an additional new MD run matching the functional reference state is created (i.e. matching the MD validation run). The learning performance of this run is compared to the MD validation run using a canonical correlation analysis conducted using all selected learners (i.e. the stacked model) across both space and time (i.e. fluctuations backbone atoms of individual amino acids over subdivided time intervals). Any sequence dependent or ‘functionally conserved’ dynamics can be recognized through a significant canonical correlation in the profile of the overall learning performance along the amino acid positions for the two similar state runs. In effect, this metric defines dynamics that are functionally conserved by capturing a signal of significant self-similarity in dynamics that co-localizes to a specific part of the protein backbone.

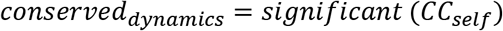

Significantly conserved regions calculated within a user defined sliding window (default value =20 residues with cutoff of p<0.01) can be plotted upon the positional local correlational value profile (i.e. R value) and also can be mapped to the reference structure of the protein, colored in dark gray on a light background.

### Variant impact assessment

By extension, mutational impacts of genetic or drug class variants on the functionally conserved dynamics can be quantified by their effects that range significantly beyond that observed in the self-similar reference runs. Thus when canonical correlations of variants differ significantly from the self-correlation observed in functionally conserved regions, according to a bootstrap test, we can plot the magnitude of impact defining how the variant’s dynamics differs from the routine self-similar dynamics of the normal functioning protein. The impacts of dissimilar states caused by altered amino acid sequence or different binding partners are assessed through their local effect on the same canonical correlation identifying conserved dynamics. We introduce a metric of relative entropy relating the canonical correlations in both the self-similar and altered variant state. In essence, this is a metric of the ‘impact’ of a given genetic or drug class variant within the context of normal functioning dynamics. For example, when trained on a natural binding interaction (e.g. DROIDS analysis comparing a DNA binding protein in its bound and unbound states), novel MD simulations with a variety of amino acid replacements can be deployed to see whether the learners can still recognize the functional dynamics in the mutant forms. In this case, functionally tolerated mutations will result in functionally conserved dynamics that do not vary outside of ± 3 standard deviation bounds of the self-similar runs, whereas functionally intolerant mutations will result in significant deviations from self-similarity of motion. An overall impact of a genetic and/or drug class binding variant on the conserved dynamic regions is calculated by

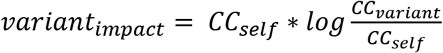

Comparative plots of local variant impacts outside of the 3 standard deviation bound determined by the validation run are generated within a user defined sliding window. Thus, this variant impact metric is designed to identify variant regions with dynamics that potentially alter disrupt conserved dynamic features of the normal functioning protein system.

### Four example applications (case studies)

To demonstrate the performance and utility of DROIDS 3.0 + maxDemon 1.0, we ran the following four comparative case studies using the PDB IDs mentioned below. Bound and unbound files were created by deleting binding partners in UCSF Chimera and resaving PDBs (e.g. 3t0z_bound.pdb, 3t0z_unbound and 3t0z_ligand). Each MD run ensemble consisted of 200 production runs at 0.5ns each explicitly solvated in a size 12 octahedral water box using TIP3P solvent model with constant temperature under an Anderson thermostat. The models were charge neutralized with both Na+ and Cl-ions. The heating and equilibration runs prior to production were 0.3ns and 10ns respectively. Prior to heating 2000 steps of energy minimization were also performed. All seven available machine learning classifiers were trained on the functional MD ensembles and deployed upon new 5 ns production runs for each variant analyzed.

Case study 1 (figure 2) – PDB ID = 1ubq – to analyze self-stability and effect of temperature shift in ubiquitin

**Figure 2.**
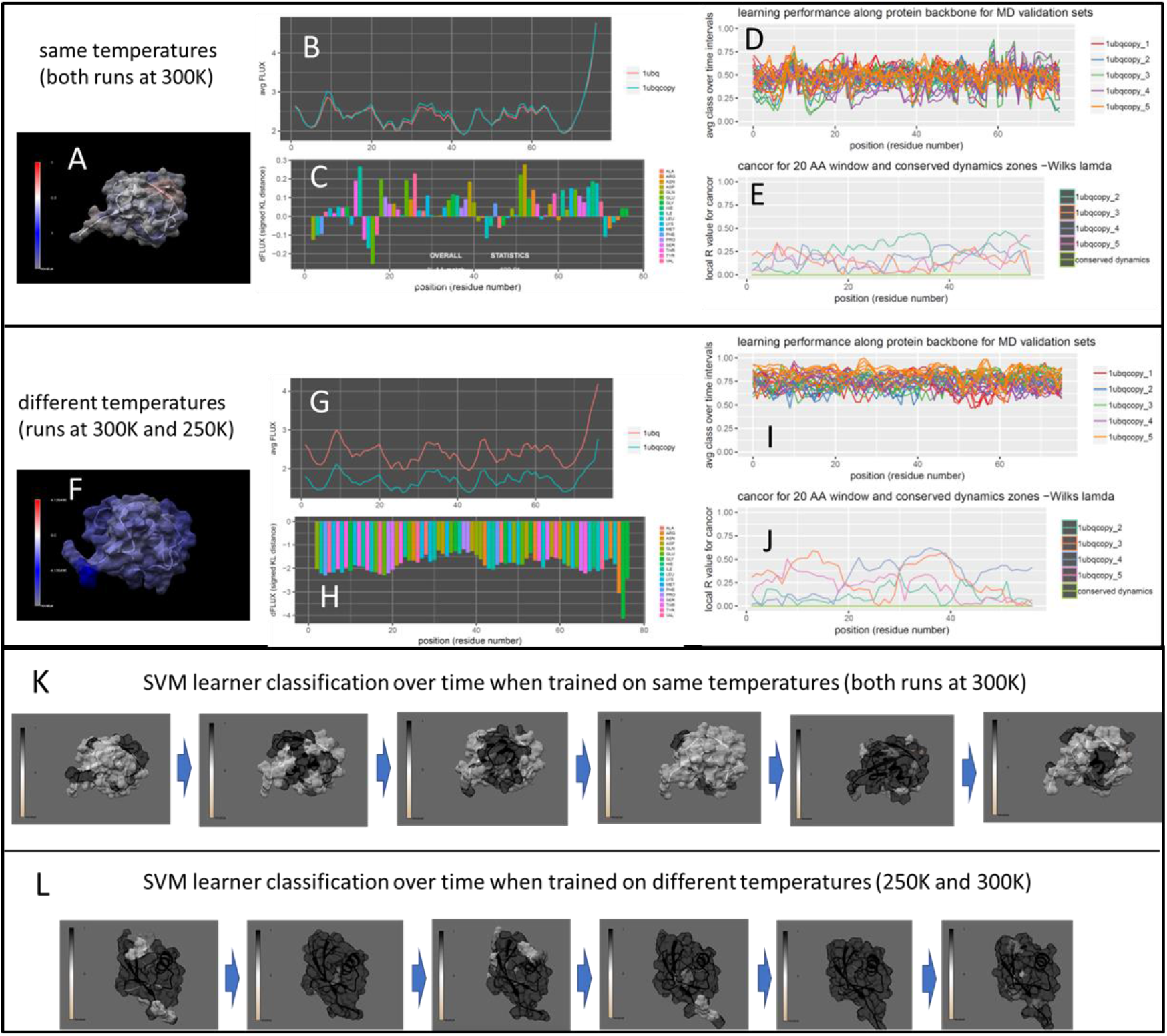
Analysis of environmental temperature change on non-functional ubiquitin dynamics. DROIDS image and analysis of random ubiquitin dynamics compared at the same (A-E) and different (F-J) temperatures. Note: blue color quantifies damped rmsf at temperature lowered by 50K. Note that performance is much higher when a temperature difference is modeled (D and I resp), however, as expected, neither comparison offers the machine learners a sequence-dependent profile by which to establish a signal of conserved dynamics (E or J). The learner classifications for the best performing learner in this case (quadratic discriminant function: QDA) is shown imaged on the ubiquitin structure over time in both the (K) random dynamics and (L) temperature dampened dynamics. (Movies of this can be observed in supplemental file A)

Case study 2 (figure 3) – PDB ID = 2oob – to analyze functional binding of ubiquitin to ubiquitin ligase and impacts of several tolerance pre-classified genetic variants

**Figure 3.**
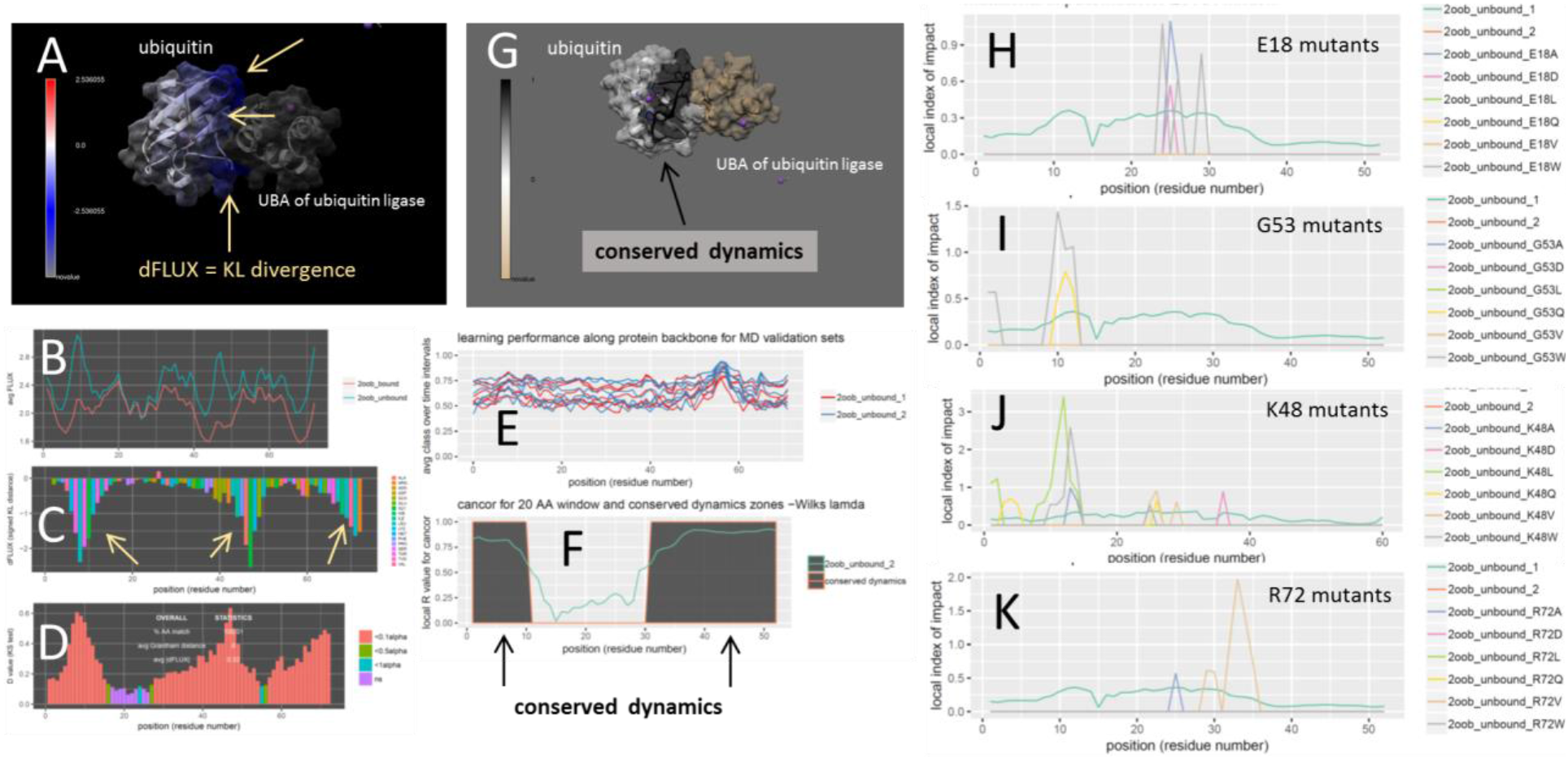
Analysis of mutational impact and tolerance on functional ubiquitin dynamics. (A) DROIDS image and analysis of ubiquitin bound to the ubiquitin associated binding domain (UBA) of ubiquitin ligase. Note: blue color quantifies damped rmsf at binding interface. (i.e. negative dFLUX) also by the (B) respective rmsf profiles of bound and unbound training states and (C) the KL divergence or dFLUX profile colored by residue. Arrows indicate most prominent dampening of rmsf near loops at THR 9, ALA 46 and C terminus. (D) Significant differences in these rmsf profiles is determined by multiple-test corrected two sample KS test. (E) Local learning performance of each machine learning method in self-similar testing runs are shown color-coded by run and regions of functionally conserved dynamics, determined via significant local canonical correlation are shown in dark gray in both (F) traditional N to C terminal plot as well as (G) structural image. The mutational impacts of 24 genetic variants (H-K: six variants at each or four sites) are shown all demonstrating lack of impact in functionally conserved regions of the binding interaction.

Case study 3 (figure 4) – PDB ID = 1cdw – to analyze functional binding of TATA binding protein to DNA and impacts of several genetic variants

**Figure 4.**
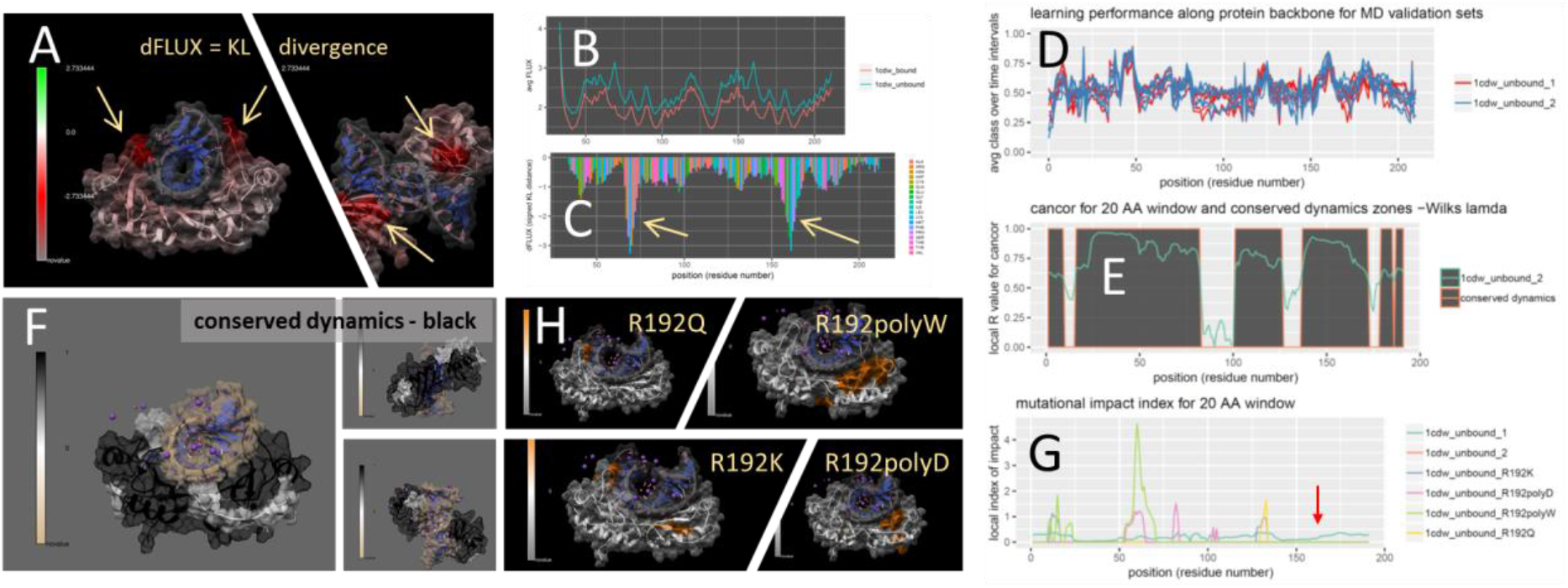
Analysis of mutational impact and tolerance on DNA binding in Tata Binding Protein (TBP). DROIDS image and analysis of TBP in DNA-bound and unbound states showing (A) colored TBP structure, (B) respective rmsf profiles and (C) KL divergence (dFLUX) plot. Note: arrows indicate functional binding loops in the DNA minor groove red color indicates dampened rmsf. maxDemon analysis (D-E) identifying conserved dynamics supporting both minor groove binding loops and (F) connecting them through the central region of the beta sheet in the main body of TBP closest to the DNA. Mutational impacts of 4 genetic variants with increasing impact one of the functional loops are also shown (G) plotted and (H) on the TBP structure. They are R192K, R192D, R192Q and polyW centered at R192 in 1cdw.pdb and and position 161 (red arrow) in plots (Note: 31 position offset is due to DNA in the original file).

Case study 4 (figure 5) – PDB ID = 3t0z – to analyze functional ATP binding in Hsp90 and subsequent impacts of six inhibitor drug variants

**Figure 5.**
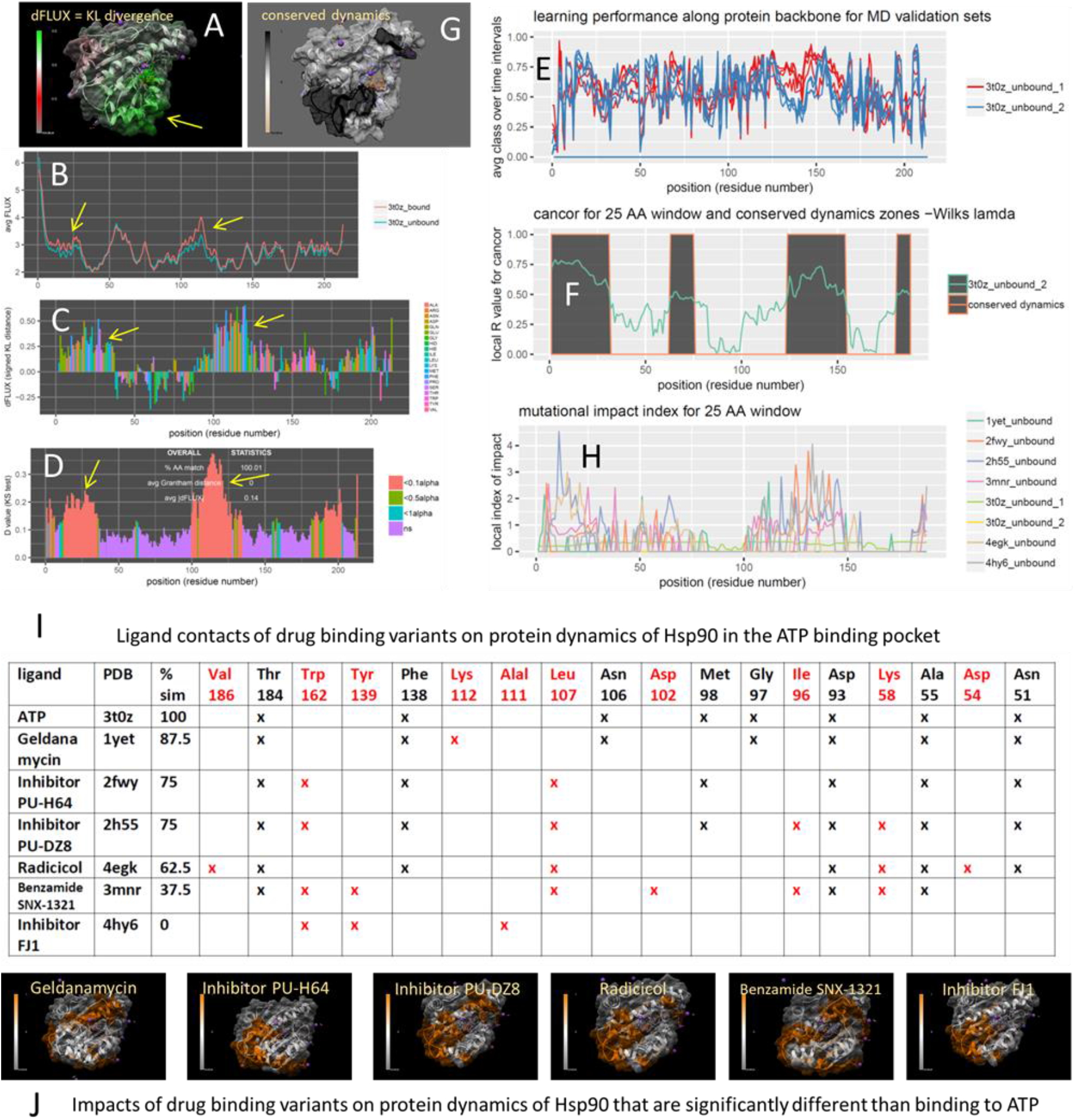
Analysis of drug class variant binding in the ATP-binding domain of Hsp90. DROIDS image and analysis of Hsp90 in ATP-bound and unbound states showing (A) colored Hsp90 structure, (B) respective rmsf profiles and (C) KL divergence (dFLUX) plot and (D) significant differences in dynamics determined via the KS test. Note: arrows and green color indicate regions where rmsf is amplified in response to ATP binding. maxDemon analysis (E-G) identifying conserved dynamics connecting the ATP binding pocket and region of amplified rmsf. (H) Mutational impacts of 6 drug class variants targeting the ATP binding pocket of Hsp90 are plotted and (I) ordered by number of differences in structural contacts within the binding pocket. (J) Mutational impacts of these variants are demonstrated to predominantly impact the functionally conserved region of amplified rmsf thus mimicking the dynamic effect of functional ATP binding.

### Improvements and upgrades over previous versions

To enhance the user experience and scientific utility, DROIDS v3.0 offers many new features beyond earlier major release versions 1.2 and 2.0. These are summarized below.

- New GUI organization directs users to specific comparative tasks/applications in Table 1
- A new control file builder for managing path dependencies in Linux is included
- Amber16/18 support has been beta tested and is defined via paths.ctl file
- Single or dual GPU user options are available for faster analyses
- Automated structure prep (dry and reduce) via pdb4amber is now included in the GUI. The ‘reduce’ variable is optional allowing users to either setup their own protonation states ahead of DROIDS, or simply allow DROIDS to hydrogenate the input structures entirely.
- Program/package dependency installer script named ‘DROIDSinstaller.pl’ is included. It will lead users through all dependencies required after a fresh Linux build, including CUDA libraries and tools required for Nvidia GPU accelerated Amber in the Linux environment
- KL divergence (= relative entropy) definition of dFLUX is now included as an option providing a richer color mapping of dFLUX in images and movies than the simple averaging algorithm offered in earlier DROIDS versions
- Binding interaction analysis for both protein-DNA and protein-ligand systems is now offered with dedicated GUI for these comparisons. Protein-ligand system setup includes QMMM preprocessing in Antechamber and SQM.
- LeAP control files for explicit solvent runs are now presented for advanced user modifications (e.g. changing ion concentration, water model, water box dimension of volume).
- Dedicated GUI allowing genetic mutation placement (on DNA or AA) are included for setting up variants to analyze
- Self-stability and temperature shift analysis has its own dedicated GUI, allowing users to copy the input pdb file to compare MD ensembles generated on identical structures at the same of at different temperatures
- MaxDemon 1.0 – machine learning based detection of functionally conserved dynamic regions
- MaxDemon 1.0 – machine learning based impact assessment of variants (genetic, structural or binding)
- Dynamic visualization and movie rendering of machine learning classification performance
- Virtual reality and ChimeraX compatibility is also supported (additional information and download code can be found here https://cxtoolshed.rbvi.ucsf.edu/apps/moleculardynamicsviewer https://github.com/kdiller713/ChimeraX_MolecularDynamicViewer

### Basic implementation of DROIDS and maxDemon

The first step of any DROIDS analysis is to find or create two homologous PDB file format structures that represent the query and reference functional states of the protein system under investigation. Typically, these would represent the same protein in a bound vs. unbound state, or in a mutant vs. wildtype state. If the protein is interacting with a small ligand, and additional ‘ligand only’ PDB file should also be created for subsequent quantum mechanical optimization and preparation by Ambertools antechamber program. These files should be placed within the DROIDS download folder. Upon implementation via the command ‘perl DROIDS.pl’ launched from terminal within the DROIDS folder, the DROIDS graphical user interface (GUI) will help the user write a control file for required working path directories on their system (first use only) and then proceeds to a main GUI outlining the various types of comparisons that can be generated (as detailed in Table 1) and the number of GPU available on the system. The next step is provides a user-friendly GUI to control and schedule Amber16/18 GPU-accelerated MD simulation to generate ensembles of short MD runs representing two functional protein states wanting to be compared. These functional comparisons are not limited, but would typically entail the impact of mutation (comparing dynamics before and after one or more amino acid replacements), the impact of an environmental change (comparing two states of temperature of solvent set up), or the impact of a molecular interaction (comparing bound to an unbound state). The DROIDS GUI will lead users through the building of a structural alignment file using UCSF Chimera’s MatchMaker and Match-Align tools. This will be needed later by the graphics components of DROIDS to make sure that only homologous regions of structures are being compared and analyzed. In this application, where the user is primarily interested in genetic or drug class variant impacts on an interactive signaling function, the typical training ensembles generated by DROIDS for further analysis with maxDemon should represent the normal binding function of the wild-type protein and therefore the bound vs unbound comparison would typically be used. A PDB file of the bound state can be the starting point and an unbound PDB model can be saved after deleting chains in the original file. If a small molecule ligand interaction is under study and requires application of an additional force fields such as GAFF, than an additional file representing only the ligand should also be generated and saved for preparation with antechamber software prior to building the solvent models using teLeAP. The GUI will pop open the .bat files that control more details of the simulation setup allowing advanced users to write more lines into the teLeAP modeling prep (e.g. to alter the water box dimensions, the water model itself, or to add additional ions beyond simple charge neutralization). The user should read all warnings provided to the terminal at this stage by the Amber software. Our GUI script will also double check the sizes of the files generated at this stage and will supply a warning if teLeap failed altogether to set up the complete model system for simulation. Upon successful setup the user can launch all the MD runs from the GUI. The requested jobs are automatically scheduled to each GPU one at a time by our software. When finished, the user can easily generate rmsf data by using the GUI to setup and launch cpptraj software provided in Ambertools. Thus the total process from file preparation, MD production and post-processing for DROIDS analysis by simply working down the buttons on each GUI from top to bottom and subsequently following the directions on the main terminal. After MD simulation and post-processing, DROIDS will take users to a second GUI for generating R plots and analyses for statistically comparing the dynamics, and then to a third GIU for visualization and movie generation. We refer users to our user manual and previous publication for more details. This third GUI has buttons to optionally launch our new machine learning application maxDemon if users wish to go beyond simple comparative protein dynamics and investigate novel simulations utilizing the DROIDS MD ensembles as a training set for subsequent machine learning.

More detailed instructions to users are included with our DROIDS 3.0+maxDemon 1.0 user manual available in the GitHub repository.

### The main repository for DROIDS 3.0 and maxDemon 1.0 can be found here. Please follow the link to “Releases” and download the latest release as .tar.gz or .zip file

https://github.com/gbabbitt/DROIDS-3.0-comparative-protein-dynamics and DOI: 10.5281/zenodo.3358976 concurrent with this publication https://zenodo.org/record/3358976#.XURVkOhKiiM

We also post various videos of examples using DROIDS, video tutorials, and ongoing projects here https://www.youtube.com/channel/UCJTBqGq01pBCMDQikn566Kw

## Results and Discussion

To demonstrate the variety of comparative analyses that can be addressed with the new release of DROIDS 3.0 and maxDemon 1.0, we chose four different case studies of comparative protein dynamics. These included (A) an analysis of self-stability and temperature effects in single ubiquitin structure, (B) a functional genetic variant analysis of ubiquitin and ubiquitin ligase binding interaction, (C) a functional genetic variant analysis of DNA binding in TATA binding protein, and (D) a drug class variant analysis of compounds targeting the Bergerat ATP binding region of Hsp90 heat shock protein.

### Machine learning analysis of impacts due to simple environmental temperature shift

We first ran a null comparison as a ‘sanity check’ by running a query and reference ubiquitin (11) MD at the same temperatures (both 300K) and same solvent conditions. The DROIDS analysis (Figure 2A-C) showed identical atom fluctuation profiles along the backbone and a random dFLUX profile indicative of nonsignificant differences due to small random local thermal differences in the training sets. The machine learning classification plots on new MD runs vary randomly around 0.5 reflecting the fact that the learning algorithms effectively had no features to train on (Figure 2D). As expected, no significantly conserved dynamics were identified either (Figure 2E). By contrast, a protein dynamic comparison run with a 50K temperature difference (Figure 2 F-H) shows a much higher machine learner performance upon deployment (i.e. 70-80% successful classification – Figure 2I). Because environmental temperature shifts are not expected to reflect evolutionary conserved dynamics (i.e. are not position dependent in their effect), they also subsequently do not result in significant canonical correlations in the learning profiles (Figure J). Representative time slices of the positional classifications in each of these experiments are shown in K and L resp and indicate that our machine learning is capable of extracting and identifying simple differences in dynamics due to temperature. Another interesting observation here was the slightly higher learning performance of the simpler machine learning methods QDA and LDA over others at all sites in the temperature shifted example. We interpret this to be related to the fact that underlying rmsf distributions are probably Gaussian, a critical assumption of these two models, with unequal variances caused by steric hindrances on the backbone. This would predict that QDA might outperform other learners in this situation and it appears that it does. We note that where more complex functional dynamics are concerned, the more sophisticated learning methods such as support vector machine and adaboost often perform slightly better than others. However, we also note that these performance differences are usually quite small and that all learning methods generally come to similar local conclusions about functional dynamics. We examine machine learning performance regarding more functional binding dynamics in ubiquitin.

### Machine learning analysis of impacts of genetic variants on a functional protein binding interaction

To examine functional dynamics in ubiquitin, we conducted a DROIDS analysis comparing its two functional states, bound and unbound to the ubiquitin associated binding (UBA) domain of ubiquitin ligase (12)(Figure 3A-D). This binding domain is highly conserved among the many other proteins that interact directly with ubiquitin. The binding interaction greatly reduces the atom fluctuation in ubiquitin at 3 characteristic positions, two loop structures centered at LEU 8 and ALA 46 and a portion of beta sheet at the C terminus (Figure 3C). These three regions also drive significant differences in dynamics across the whole protein. In novel self-similar MD runs on the bound state, we successfully detect significant canonical correlations indicating conserved dynamics in these three regions with a broad expanse in conserved dynamics (Figure 3E and F) across the UBA region (Figure 3G). We tested a set of 24 mutations that included sites with the most and least tolerated effects on growth rate *in vivo* in yeast according to a study by Roscoe et al. (13). In this study, nearly all mutations at E18 and G53 are tolerated while nearly all mutations at K48 and R72 are not. Ultimately, the causes of tolerance in these variants are not known, and do not necessarily invoke functional problems in dynamics. However, the impacts that we did observed in simulation were on average twice as strong in the intolerant backgrounds when compared to the mutation tolerant backgrounds. And, while we did not see large differences in the number of mutational impacts on dynamics between tolerated and non-tolerated mutant groups, the 24 mutations analyzed all show a general trend of dynamic impact falling outside of most of the functional binding region (Figure 3H-K), suggesting that ubiquitin may have evolved a tertiary structure that allosterically translates dynamic impacts to less functional regions of the protein. Some interesting exceptions to this rule were demonstrated by the very large impacts of K48L, K48W and R72D, centered squarely in the functionally conserved binding regions of ubiquitin, and would obviously heavily disrupt electrostatic charge interactions there as well.

### Machine learning analysis of impacts of genetic variants on DNA binding interaction

TATA binding protein (TBP) is a general transcription factor that binds DNA upstream in most highly regulated eukaryotic gene promoter regions (14). While relatively small, it is a mechanically dynamic protein with a C-clamp like structure that highly distorts the rigid DNA double helix by inserting four phenylalanine side-chains between base pairs. It is thought that this bending allows TBP to be more rapidly released from the TATA element, as opposed to TATA-less promoters, subsequently allowing more highly controlled regulatory responses in TATA box genes (15). Due to its obvious symmetry and ability to impart large forces during binding, we thought that it would represent a good candidate for comparison of its dynamics during its binding interaction with DNA. We conducted a DROIDS analysis comparing human TBP (16) in its functionally bound and unbound states (Figure 4A-C). TBP exhibits a characteristic large signature of dampening of atom fluctuation throughout its entire structure with most pronounced effects in two loop regions that interact with the minor groove of DNA (arrows in Figure 4A and 4C). Canonical correlations in new self-similar MD runs marking increased performance in classification were observed in these regions (Figure 4D) along with corresponding regions of conserved dynamics identified by significant Wilk’s lamda (Figure 4E). Conserved dynamics from these loop areas are connected through the chains in the beta sheet region of TBP spanning the DNA major groove contact. Mutational impacts of four variants affecting the binding loop most proximal to the C terminal exhibited followed our expectation of increasing impact ordering from R192Q, R192K, R192polyD, and R192polyW (Figure 4G and 4H). The polyD and polyW mutations incorporated 5 sequential ASP or TRP residues centered at R192, both causing the loop region to become more rigid (causing increased negative dFLUX). We expected the strong functional binding affect observed across nearly all residues in this system would make it relatively highly tolerant to single amino acid substitutions, even when located in the most functional binding loop. In accordance with our expectations, we found the most impactful multiple mutation (i.e. R192polyW) significantly affected the dynamics of nearly 6 times more local residues than the least impactful single substitution (i.e. R192Q).

### Machine learning analysis of impacts of drug class variants targeting the ATP binding region of Hsp90

In contrast to TBP, we wanted to use our method to examine a small molecule binding interaction in a protein with potentially more complex impacts on molecular dynamics. Hsp90 is a well-known chaperone protein that assists the folding of many proteins and thereby mitigating many environmental stresses in the cell. Hsp90 even capacitates the evolutionary process by allowing potential phenotypic variation exhibited under stress to be hidden from natural selection until needed in response to environmental change (17). Hsp90 contains a highly conserved N-terminal domain where ATP binding and activation occurs. The binding of ATP physically changes motions in this region creating a ‘lid’ that closed during ATP binding and open when conversion to ADP occurs. Due to the role of Hsp90 in stress mitigation in most tumors, it is a common drug target for ATP inhibitors in many cancer therapies (18, 19). The amino acid residues that interact with ATP in this region are well known and the inhibitor geldanamycin is known to mimic nearly all the local ATP contacts as well (20). Other more modern inhibitors interact with the ATP binding pocket quite differently (19, 21, 22), so we thought that this system would be a good candidate for comparative analysis of drug class variants with our software.

We conducted a DROIDS analysis comparing the dynamics of Hsp90 chaperone, a common drug target for inhibitors in many cancer therapies, in both its ATP bound and unbound states. The binding of ATP was discovered to significantly destabilize three co-localized alpha helical regions of the protein adjacent to and extending from the ATP binding site (Figure 5A-D). MaxDemon analysis confirmed the dynamics of this region to be highly conserved in new MD runs (Figure 5D-G). We also analyzed the impacts of the six drug class variants targeting the ATP site (20, 22, 21, 23, 24), but interacting differently with residues in this region (Figure 5H). The contacts in the ATP binding site are shown in Figure 5I. While the localized patterns of impacts of the drug variants were all quite similar to ATP (Figure 5J), the drug variants that most closely mimicked the contacts of ATP (i.e. geldanamycin) had far less impact on conserved dynamics than variants that interacted very differently with the binding pocket (i.e. benzamide SNX1321 and inhibitor FJ1(Figure H-I). We feel that this finding demonstrates not only demonstrates the potential of our method/software quite well, but it also demonstrates that while it is important to be able to target a druggable protein binding site (25), researchers should also consider how these various small molecules might alter, or fail to alter, the natural dynamics of the system. In situations where a drug might too closely mimic the dynamic effects of a natural activator like ATP, a hyperactivation response might occur in non-tumor cells leading to secondary cancer (26–28). Alternatively, other situations may require drug targeting that does not alter the natural dynamic behavior too much, potentially activating proteolytic systems in the cell. Our software allows more detailed investigations of these potential dynamic impacts of drug class variants.

### Conclusion

We provide a well demonstrated method and user-friendly software pipeline for conducting statistically sound comparative studies of large ensembles of comparative protein dynamics. The method/software also now provides machine learning based extrapolations of effects on novel MD simulations representing various functional variants of interest to the user. While there currently is at least one other software allowing users to connect sequence-based evolutionary metrics to protein dynamics (29), our method/software is unique in that regions of functional conservation are identified by analyzing self-similar features of dynamics themselves rather than relying upon marrying dynamics analysis to traditional static sequence-based approaches, which do not necessarily assume that a conserved function region has a strong dynamic component. By providing a systematic way of comparing protein dynamics at single residue resolution, our method/software provides an important step beyond traditional sequence-based bioinformatics, allowing investigators to gain a much more biophysically-grounded view of functional and evolutionary change. Another advantage to our method/software is that our functional impacts (i.e. mutational tolerance) are defined solely within the context of protein dynamic system being simulated. This provides a much deeper look into protein specific function than current genomic and proteomic database methods of predicting mutational tolerance (30, 31) currently allow. As GPU technology continues to advance at a rapid pace over the next few years, our method/software may have profound potential application to the development of precision and personalized medicine, where understanding the detailed interaction between genetic and drug class variants within the context of specific protein dynamic systems will be greatly needed.

## Supporting Material

The main repository for DROIDS 3.0 and maxDemon 1.0 can be found at the GitHub repository link below. Please follow the link to “Releases” and download the latest release as .tar.gz or .zip file https://github.com/gbabbitt/DROIDS-3.0-comparative-protein-dynamics We also post various videos of examples using DROIDS, video tutorials, and ongoing projects here https://www.youtube.com/channel/UCJTBqGq01pBCMDQikn566Kw A ‘live version’ of the figures in this manuscript is also available on our YouTube channel.

## Author Contribution

GAB and EPF conceived the project and method. All authors contributed to the code base. GAB and LEA worked on beta testing and debugging.

## Acknowledgements

We acknowledge the lab of Dr. Andre O. Hudson for beta testing our software during its early development. We also acknowledge Dr. Daniel Wysocki (Astrophysics program-CCRG at Rochester Institute of Technology) for very helpful suggestions regarding early methodological discussions. We also acknowledge Dr. Miranda Lynch and her colleagues at the Hauptman-Woodward Medical Research Institute for many helpful comments.

